# Detection of Frustration-related Operant Behavior in Rats via Machine Learning Methods

**DOI:** 10.64898/2026.08.26.747319

**Authors:** Jiefei Wang, Anirudh Babu, Bach Nguyen, Yorkiris Marmol Contreras, Poonam Shah, Isabella Carmona Ramirez, Thomas A. Green

## Abstract

Despite its strong link to neuropsychiatric conditions, frustration remains critically understudied in humans and animals alike. Therefore, there is an urgent need to develop tools to understand and therapeutically target frustration-related functions. Interestingly, humans and rats respond similarly during frustrative nonreward by increasing barpress durations. We previously validated barpress duration in rat operant tasks as a reliable measure of frustration-related behavior; however, it is well-known that in addition to duration of responding, emotional states such as frustration alter other aspects of responding such as force of pressing. One-dimensional, static measures such as maximum force could miss rich information contained within operant data. Thus, the objective of this study is to apply machine learning (ML) to force/time profiles to discriminate frustration-related barpresses from non-frustration-related barpresses. Results showed an AUROC for FR1 (i.e., non-frustrated) vs. extinction (frustrated condition) for individual barpresses of 0.65 that improved to 0.84 with a chunk size of 10. The model generalized well to progressive ratio responding, a different kind of frustration procedure. We conclude that force/time profiling does provide utility beyond one-dimensional measures of duration or force separately, meaning that we can indeed infer the internal state of frustration from behavior using ML techniques. Importantly, this project will also serve as proof-of-concept for applying ML to predict other internal states from barpress data.

## 1 Introduction

Frustration is a complex affective state that emerges from the interplay between external contingencies and internal motivation[1–3]. In the context of animal behavior, understanding frustration can provide insights into how animals respond to their environment and eventually help in understanding the corresponding neural mechanisms in humans. However, determining frustration in animal behavioral models is challenging because animals cannot directly communicate their affective states. As a result, researchers must rely on animal behavior to infer their affective states[1, 2]. Prior work used speed running down a runway or amount of sucrose consumed to infer frustration state[4, 5]. However, these techniques did not provide sensitive timecourse data (i.e., changes in frustration across a session). Our recent work showed that barpress durations can be used as a proxy measure of frustration state throughout the session for normal operant responding for sucrose or drug in rats[6–8] and is useful in psychopharmacological studies[9]. All manner of frustrative nonreward (FNR) increases barpress durations, including reward downshift, reward omission, or increased price of a reward. These results showed that emotional state can indeed be inferred from the topography of a barpress, in this case durations

Humans, increase force in addition to durations when frustrated[10, 11]. However, we have previously found no reliable changes in simple max force calculations (unpublished data). The question at hand is if the force/time profile can add additional useful information beyond simple duration and max force calculations. We find here that the force/time profile of barpresses does contain a rich source of information. We hypothesized that ML applied to these profile data would provide a more accurate measure of frustration state than using durations alone. However, it remained unclear which profile data, or so-called features, are most informative for predicting the frustration state.

To answer this question, in this study, we aimed to 1) derive quantitative features from barpress data, 2) Build a machine learning model to predict frustration state and evaluate the contributions of individual features to model performance, and 3) assess whether the model generalizes to different behavioral scenarios that also induce frustration in animals. The results of this study will help determine the fundamental relationship of FNR with motivation and will have application to the preclinical study of substance use disorders as well as other neuropsychiatric conditions related to FNR.

### 2 Methods

### 2.1 Animals

Two cohorts of Sprague-Dawley rats (Envigo, Houston, TX) were obtained at 250-300 grams (22 males for each cohort), and one cohort of Long-Evans rats was bred in-house (8 males, 11 females), for a total of 63 rats (see Figure 1). Rats were maintained in a controlled environment (22 ^*°*^C; 50% relative humidity; 12 h light/dark cycle, lights on 0600 h) in a colony approved by the Association for Assessment and Accreditation of Laboratory Animal Care (AAALAC). All procedures conformed to the NIH Guide for the Care and Use of Laboratory Animals and approved by The University of Texas Medical Branch Institutional Animal Care and Use Committee.

**Figure 1:**
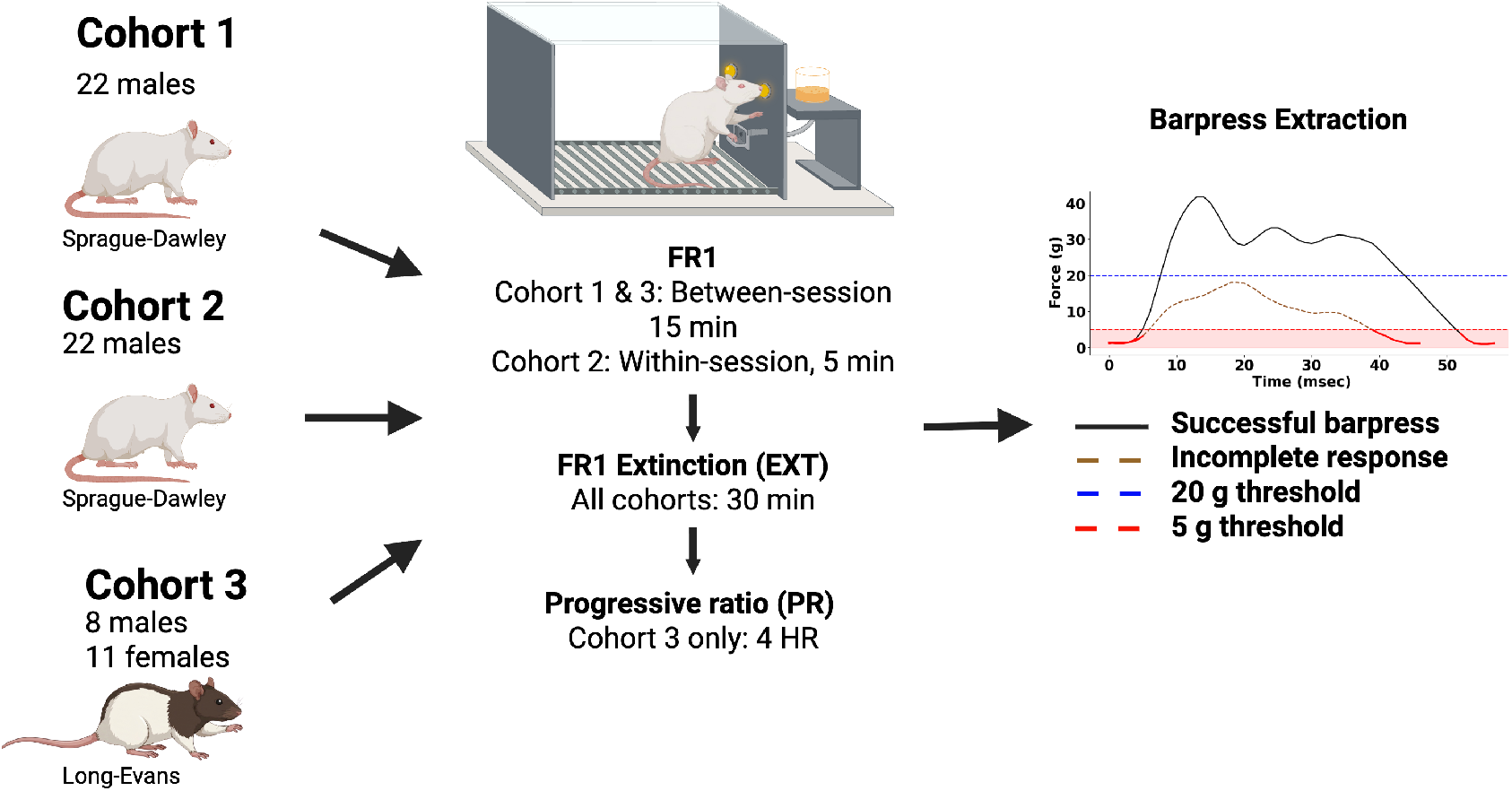
Study Diagram. Three cohorts of rats were used in the study. All rats underwent FR1 and EXT sessions with varying length. Cohort 3 underwent an addition PR session. Created with BioRender[15].

### 2.2 Sucrose training and maintenance

Rats were food regulated to 85% ad libitum body weight and then trained to press for banana-flavored sucrose pellets (45 milligrams, Bio-Serv) using standandard operant levers in a sound attenuating chamber. Upon training, operant chambers were equipped with force levers (Med-Associates, St. Albans, VT, Product Number: ENV-118M) to measure force of bar pressing. A threshold of 20 grams was required to count as a barpress and a lower threshold of 5 grams was used for the termination of barpress. When the contingency was met, two cue lights were illuminated for 5 sec and a sucrose pellet was delivered. Otherwise, a houselight was illuminated throughout the session. For active barpresses, values above 5 grams were used to extract successful barpresses. The force exerted on the lever was recorded at 100 Hz.

### 2.3 Non-frustrated responding: Fixed Ratio 1

After training, rats underwent a daily 15-min operant session at a fixed ratio 1 (FR1) schedule of reinforcement until responding stabilized. A rat was considered stable when number of reinforcements, average duration of pressing, and average maximum force of pressing showed less than 20% variability across 3 sessions. Additionally, rats must have obtained at least 30 reinforcers and a 2:1 ratio of active to inactive barpresses. Maintenance sessions were also used to stabilize responding between frustration days (maintenance). The final 15-mins period of stable FR1 responding was used in the analysis for Cohorts 1 and 3 as was the first 5-min of responding for the within-session procedure for Cohort 2.

### 2.4 Frustration: Reward omission

All cohorts underwent a reward omission task (Extinction, EXT) where successful barpresses resulted in reinforcement-associated light cues but sucrose pellets were omitted. For Cohorts 1 and 3, rats underwent EXT for a 2 hr session, with the first 30-mins used for analysis. For Cohort 2, a 30 min EXT period was made directly after the 5-min FR1 period (i.e., within-session).

### 2.5 Frustration: Progressive ratio

After maintanence FR1 sessions, Cohort 3 underwent a progressive ratio (PR) task where the contingency for each subsequent reinforcement is increased using a semi-logarithmic sequence. The first reinforcement required 1 barpress, the second reinforcement required 2 barpresses, then 4, 6, 9, 12, 15, 20 etc. The session was terminated when the animal failed to achieve a reinforcement within 60 minutes, with a maximum session length of 4-hours. These session data were used to test model generalizability to FNR other than EXT.

### 2.6 Barpress extraction

Barpress data were recorded for each session and saved as a text file along with the session metadata, and barpresses were subsequently extracted. A barpress was defined as a continuous segment of barpress data exceeding the 5 grams threshold, passing through 5-20 grams range, and peaking above 20 grams. Force samples lower than 5 grams were set to 0 gram and maximum force was capped at 400 grams. Two consecutive barpresses were separated by at least one sample of zero force. Individual barpresses were then extracted from the filtered barpress recordings. The first three barpresses from each session were excluded to allow rats to adapt their expectations.

The Med-Associates force levers occasionally produced a barpress artifact that was clearly not physiologically relevant, as the force remained steady beyond what a rat could achieve. Thus, barpresses longer than 0.5 sec with a force variation of less than 2 grams were discarded. After this filtering, we obtained a set of individual barpresses for analysis. Each barpress was assigned a label corresponding to the session type, either FR1, EXT, or PR.

### 2.7 Feature extraction

We defined a set of features from each barpress for the downstream analysis. There are three categories of features: force-based, duration-based, and distribution-based features. The force-based features include the max force, force variation rate, average first 5 values, average last 5 values, and peak sharpness; the duration-based features include the total press duration, peak number, and peak width; and the distribution-based features include skewness and kurtosis. The function has eight hyperparameters that can be adjusted to control the sensitivity of peak detection (see Supplemental Table S1 for their definition). A representative barpress force–time profile illustrating the extracted features is shown in Figure 2.

**Figure 2:**
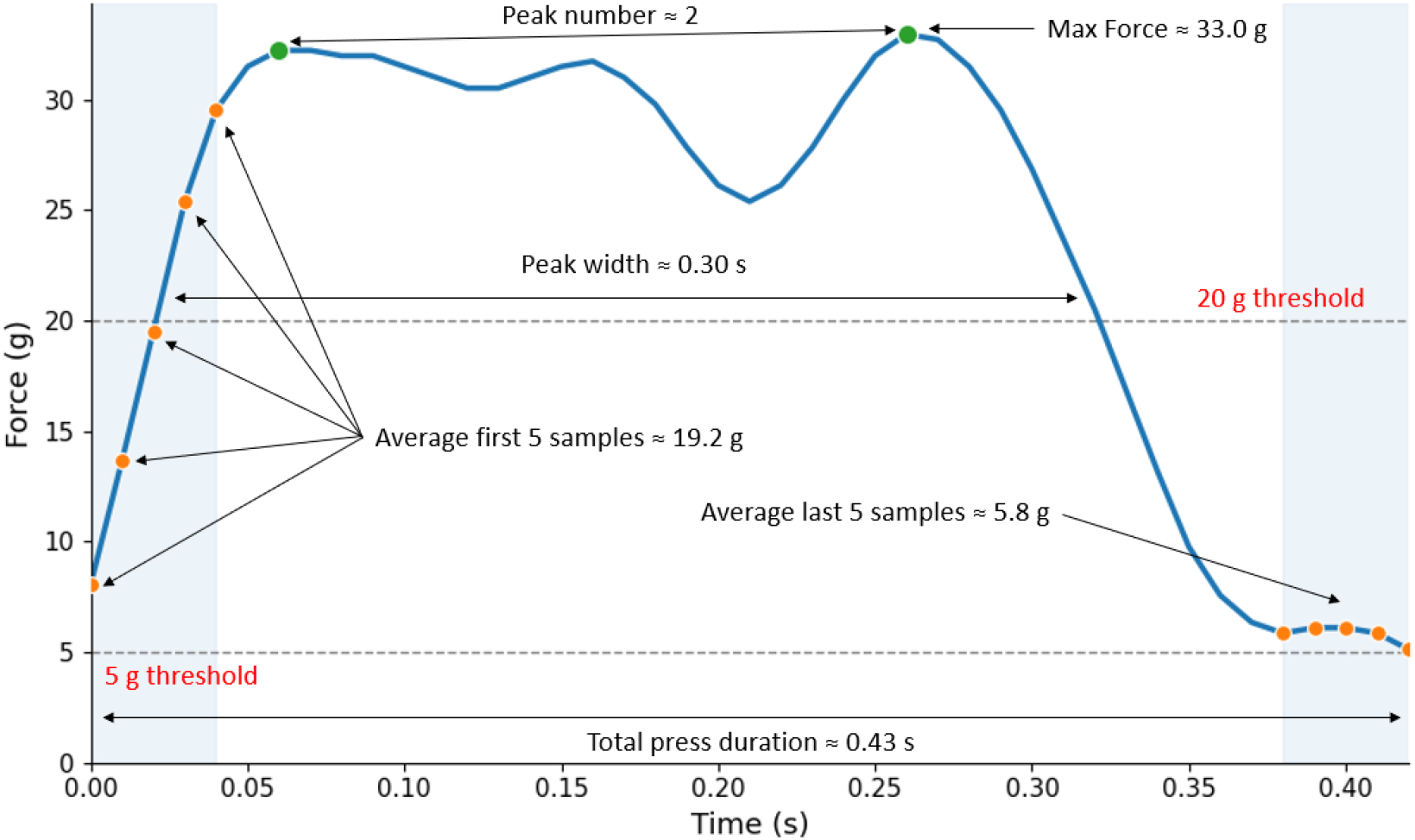
Extracted features from a representative barpress force–time profile. Illustrated features include maximum force, total press duration, peak number, peak width, and average force across the first and last five samples. Colored points mark detected peaks and the first and last five samples; dashed lines indicate the 5-g and 20-g thresholds. Peak sharpness, force variation rate, skewness, and kurtosis are not shown.

For the peak-related feature, the `find_peaks` function in the SciPy library was used to identify the peaks in each barpress. The function has eight hyperparameters that can be adjusted to control the sensitivity of peak detection. We tuned the peak hyperparameters using the training data (detailed in the section below) to extract the number of peaks, peak duration, and peak sharpness from each barpress. For barpress with multiple peaks, we aggregated values across all peaks within a barpress by computing the mean, minimum, and maximum.

### 2.8 Machine Learning Model and Training

We used a Gradient Boosting (GB) model to predict session type (FR1 vs. EXT). The inputs to the GB model included features extracted from the barpress data, along with sex, age, weight, and rat strain as rat-level covariates. We defined EXT as the positive outcome; thus, the model estimated the probability that a given barpress originated from an EXT session, which reflects a state of frustration. Hereafter, we refer to this predicted probability as the frustration score.

For model development, we performed a train/test split at the rat level[12]. Nine rats (three from each cohort, approximately 15% of the total sample) were held out for final evaluation. To account for differences in the number of barpresses across rats, we weighted each barpress by the inverse of the total number of barpresses made by that rat. Peak-detection hyperparameters used in `find_peaks` (e.g., `prominence, height,` and `distance`) were optimized using only the training data. For each candidate parameter combination, peak-related features were extracted and used to train a GB model to discriminate between FR1 and EXT sessions. Performance was evaluated using 10-fold group cross-validation with rat ID as the grouping variable, and a grid search was conducted to identify the parameter combination that maximized the mean cross-validated area under the receiver operating characteristic curve (AUC).

The search ranges for each hyperparameter are provided in Supplemental Table S2. Using the optimal peak-detection parameters, peak-related features were then extracted as described in Section 2.7.

We used correlation matrix to examine pairwise correlations among features. Features with correlations greater than 0.95 were considered highly correlated. To assess the predictive utility of individual features, we trained one-feature GB models and assessed performance with the same 10-fold group cross-validation. Performance was quantified as AUC for each fold, and each feature was summarized by the mean AUC across folds with 95% confidence intervals. Within each highly correlated feature group, a single representative feature was retained based on the performance of the one-feature GB model and feature interpretability.

Following feature selection, we trained a GB model with all retained features and rat-level covariates. We tuned the model’s hyperparameters using the same 10-fold group cross-validation framework. The optimal hyperparameter combination was selected based on the highest mean cross-validated AUC, and the final GB model was subsequently trained on the entire training dataset using these optimized settings.

### 2.9 Model Evaluation and Inference

After hyperparameters were selected, the fitted GB model was evaluated on the held-out nine test rats. Performance was summarized using AUC as the primary metric. We also evaluated model performance using accuracy, precision, recall, and F1 score. Predicted probabilities were dichotomized using the threshold that maximized the F1 score on the test data. Boxplots comparing predicted frustration scores between FR1 and EXT conditions were generated for each rat in the test set.

The contribution of the features to model predictions was evaluated using sequential permutation and SHAP values[13]. Sequential permutation is an iterative algorithm that evaluates all remaining features at each step. The feature whose permutation caused the largest AUC drop was permanently permuted, and the process was repeated until all features had been permuted. This yields an AUC degradation curve with features ranked from most to least predictive. SHAP values quantify how much each feature increases or decreases the predicted frustration score relative to the average prediction. For each feature and barpress, a positive SHAP value indicates an increase in the predicted frustration score, whereas a negative value indicates a decrease. We used SHAP beeswarm plots to visualize the association between each feature and the frustration score.

To evaluate the effect of temporal aggregation on predictive performance, we grouped consecutive barpresses into chunks and calculated the mean frustration score within each chunk. We varied the chunk size from 1 to 20 barpresses and evaluated model performance by calculating the AUC at each aggregation level. The results were visualized using a line plot.

For the PR task, we selected the optimal chunk size based on the results shown in the line plot and compared changes in frustration scores over time between the PR and FR1 tasks.

## 3 Results

### 3.1 Descriptive statistics

We collected data from 63 rats, of which 51 were male and 12 were female. The total number of recorded sessions was 186 (104 FR1 and 82 EXT, some rats have multiple sessions). After excluding 1528 barpresses with constant force, we extracted 16,118 barpresses for FR1 and 8,421 for EXT. Table 1 summarizes the extracted features between EXT and FR1 after finetuning the peak algorithm. (see Supplemental Table S3 for the hyperparameter search space and selected values for the GB model). All extracted features differed significantly between FR1 and EXT sessions, although the magnitude and direction of these differences varied across features. Barpresses from EXT sessions were associated with less maximum force, longer total press duration, and broader peaks width.

**Table 1:** Descriptive statistics for Features Between FR1 and EXT Groups.

| Feature | FR1 (Mean ± SD) | EXT (Mean ± SD) | P-value |
| --- | --- | --- | --- |
| Max Force | 46.67 ± 45.13 | 45.33 ± 35.55 | 0.011 |
| Force Variation Rate | 8.75 ± 12.75 | 7.96 ± 9.68 | <0.001 |
| Average First 5 Samples | 14.81 ± 5.97 | 15.49 ± 5.81 | <0.001 |
| Average Last 5 Samples | 14.76 ± 18.43 | 14.33 ± 14.94 | 0.049 |
| Max Peak Sharpness | 3.56 ± 6.52 | 2.98 ± 4.85 | <0.001 |
| Mean Peak Sharpness | 3.36 ± 6.34 | 2.72 ± 4.70 | <0.001 |
| Min Peak Sharpness | 3.19 ± 6.29 | 2.49 ± 4.66 | <0.001 |
| Skewness | -0.01 ± 0.66 | -0.10 ± 0.68 | <0.001 |
| Kurtosis | -0.52 ± 1.04 | -0.33 ± 1.14 | <0.001 |
| Total Press Duration | 57.41 ± 49.67 | 68.78 ± 55.99 | <0.001 |
| Peak Number | 1.32 ± 0.94 | 1.52 ± 1.08 | <0.001 |
| Max Peak Width | 21.01 ± 15.25 | 25.66 ± 20.35 | <0.001 |
| Mean Peak Width | 19.04 ± 12.66 | 21.69 ± 15.77 | <0.001 |
| Min Peak Width | 17.48 ± 12.72 | 18.53 ± 15.86 | <0.001 |

Figure 3A shows the correlation matrix among all features. Notably, there were two distinct clusters of variables. The first cluster contains force-based features (e.g., `Max Force`) and Skewness, while the second contains duration-based features (e.g., `Total Press Duration`) plus kurtosis. `Max Force` and `Force Variation Rate` exhibited strong correlations. `Max Peak Sharpness, Mean Peak Sharpness,` and `Min Peak Sharpness` also showed a strong correlation. Figure 3B shows the predictive utility of individual features. All features have an AUC below 0.6. This finding is consistent with our previous study, which found no strong predictors of the frustration state[6]. After discussion with domain experts, we retained `Max Force` and `Mean Peak Sharpness` due to their conceptual simplicity.

**Figure 3:**
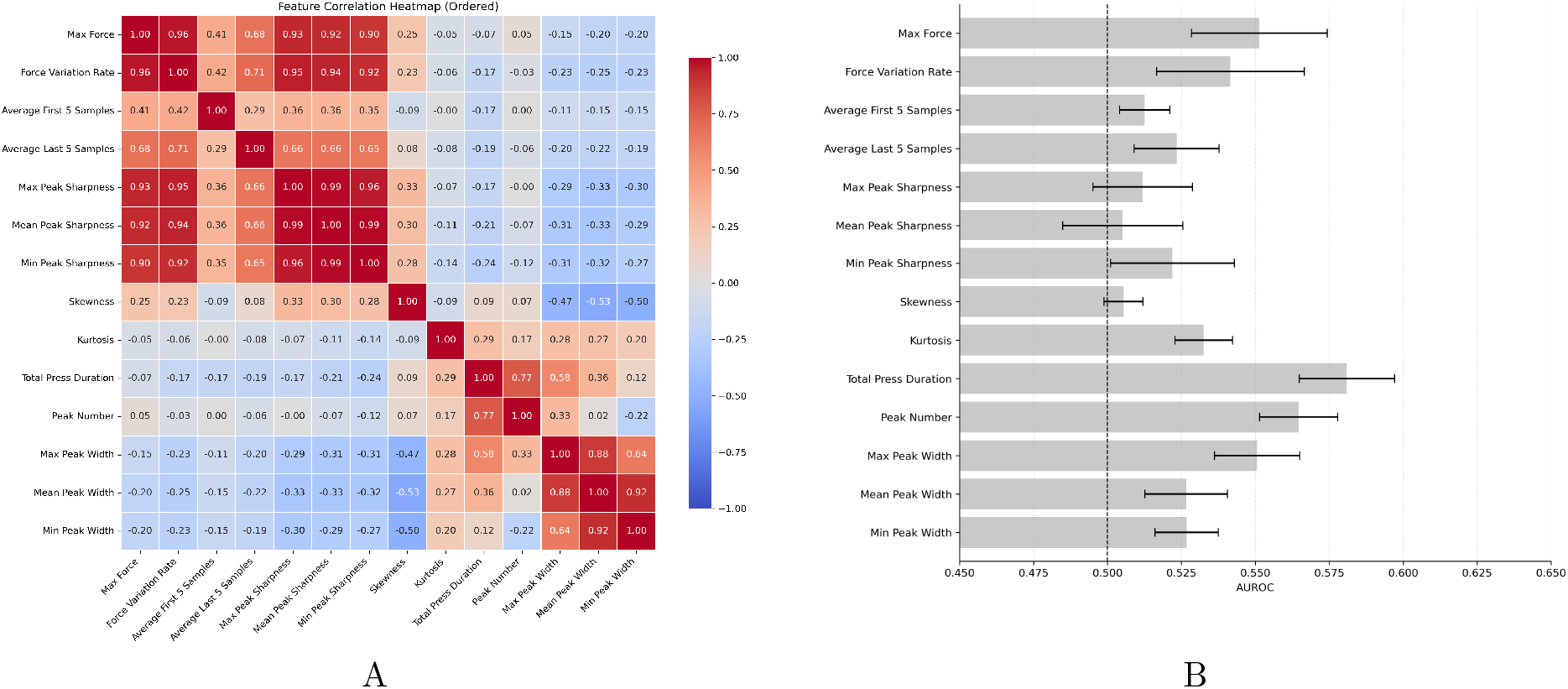
Overview of the dataset and feature evaluation results. A. Correlation structure of session data collected from rats. B. AUC for each individual feature and its *±*95% CI

### 3.2 Machine Learning

We trained GB models using the 11 barpress features and 4 rat features with different hyperparameters[14]. The best GB model achieved an AUC of 0.648 during cross-validation on the training dataset and an AUC of 0.654 on the held-out test dataset. On the test dataset, the optimal classification threshold was 0.33 with a maximum F1 score of 0.675 (precision: 0.534, recall: 0.92). At this threshold, the model correctly identified 92% of EXT barpresses. However, among the barpresses classified as EXT, approximately 47% were actually from the FR1 session.

### 3.3 Model Evaluation and Inference

The SHAP plot in Figure 4 shows the contribution of each feature for the GB model on the held-out test dataset. Features were ranked according to their mean absolute SHAP values[13]. Most features exhibited clear monotonic relationships with the predicted frustration score. For example, larger values of `Total Press Duration, Average Last 5 Samples, Max Peak Width,` and `Average First 5 Samples` were associated with increased frustration scores. In contrast, higher values of `Max Force` and `Skewness` were associated with lower frustration scores. Notably, some features exhibited U-shaped relationships with the frustration score, including `Mean Peak Sharpness` and `Peak Number,` for which high feature values were associated with both low and high frustration scores.

**Figure 4:**
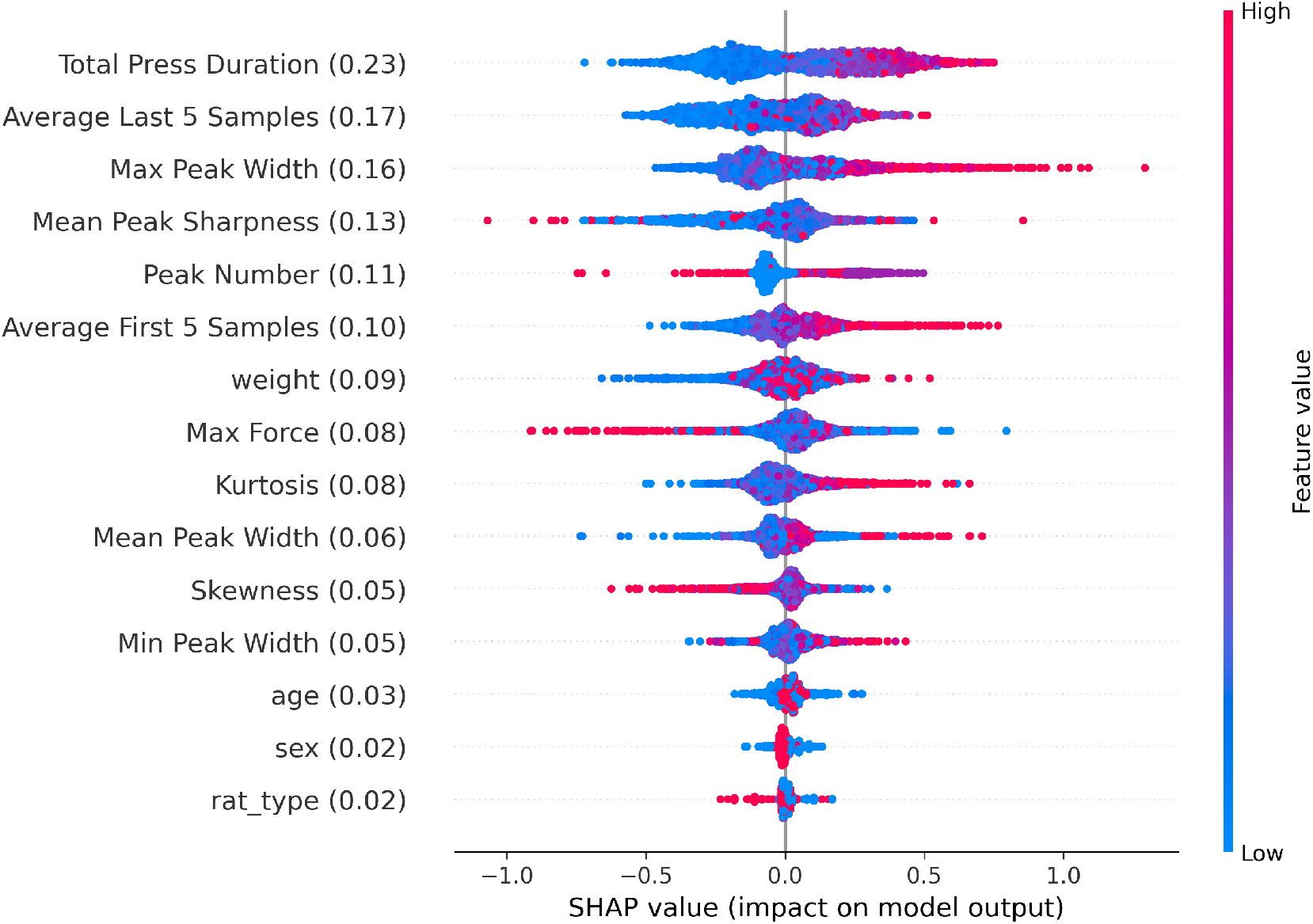
SHAP values for the GB model on the held-out test dataset. The mean absolute SHAP value for each feature is shown in parentheses after the feature name. For each feature, red color represents higher feature value and blue represents lower. Positive SHAP values indicate that the feature increases the predicted frustration score, whereas negative SHAP values indicate that it decreases the predicted frustration score.

We sequentially permute the features from the feature with most AUC drop to the feature with less AUC drop to see the degradation of the model performance in Figure 5(A). `Max Peak Width` was the frist dropped features (AUC drop: 0.037), followed by `Total Press Duration` (AUC drop: 0.0041) and `Average First 5 Samples` (AUC drop: 0.027) and `Peak Number` (AUC drop: 0.014). Together, these four features accounted for a total AUC drop of 0.111, whereas the remaining eight features contributed only an additional 0.043 AUC to the model performance (Assuming 0.5 AUC as the chance-level performance).

**Figure 5:**
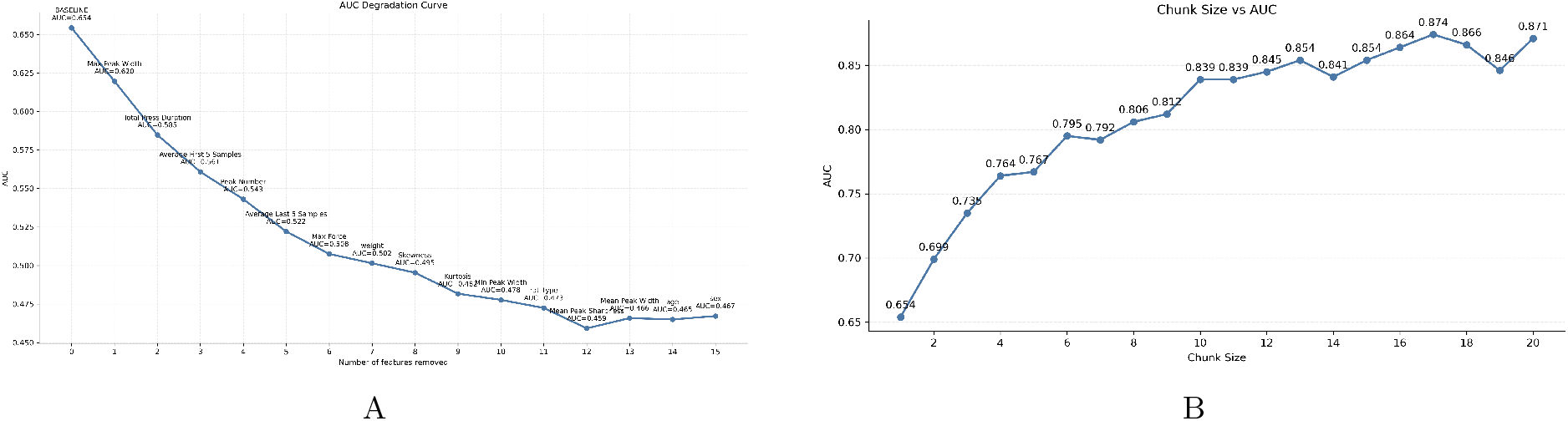
A. AUC degradation curve. B. Comparison of AUCs across chunk sizes.

To further evaluate the capacity of the GB model, we combined predictions from multiple barpresses within a temporal chunk to increase discriminative power. Figure 5B shows the relationship between chunk size and AUC. The curve exhibited an elbow-shaped pattern, with AUC increasing substantially as chunk size increased. However, the improvement diminished at larger chunk sizes. For example, the AUC reached 0.839 at a chunk size of 10 and increased only slightly to 0.871 at a chunk size of 20.

For evaluating the prediction result, the boxplot in Figure 6 shows the distribution of predicted frustration score for FR1 and EXT conditions based on the GB model in the hold-out test set. Across all rats, EXT sessions consistently show higher median predicted probabilities than FR1 sessions. FR1 distributions tend to display smaller variability compared to EXT (FR1 SD 0.146 vs EXT SD: 0.157).

**Figure 6:**
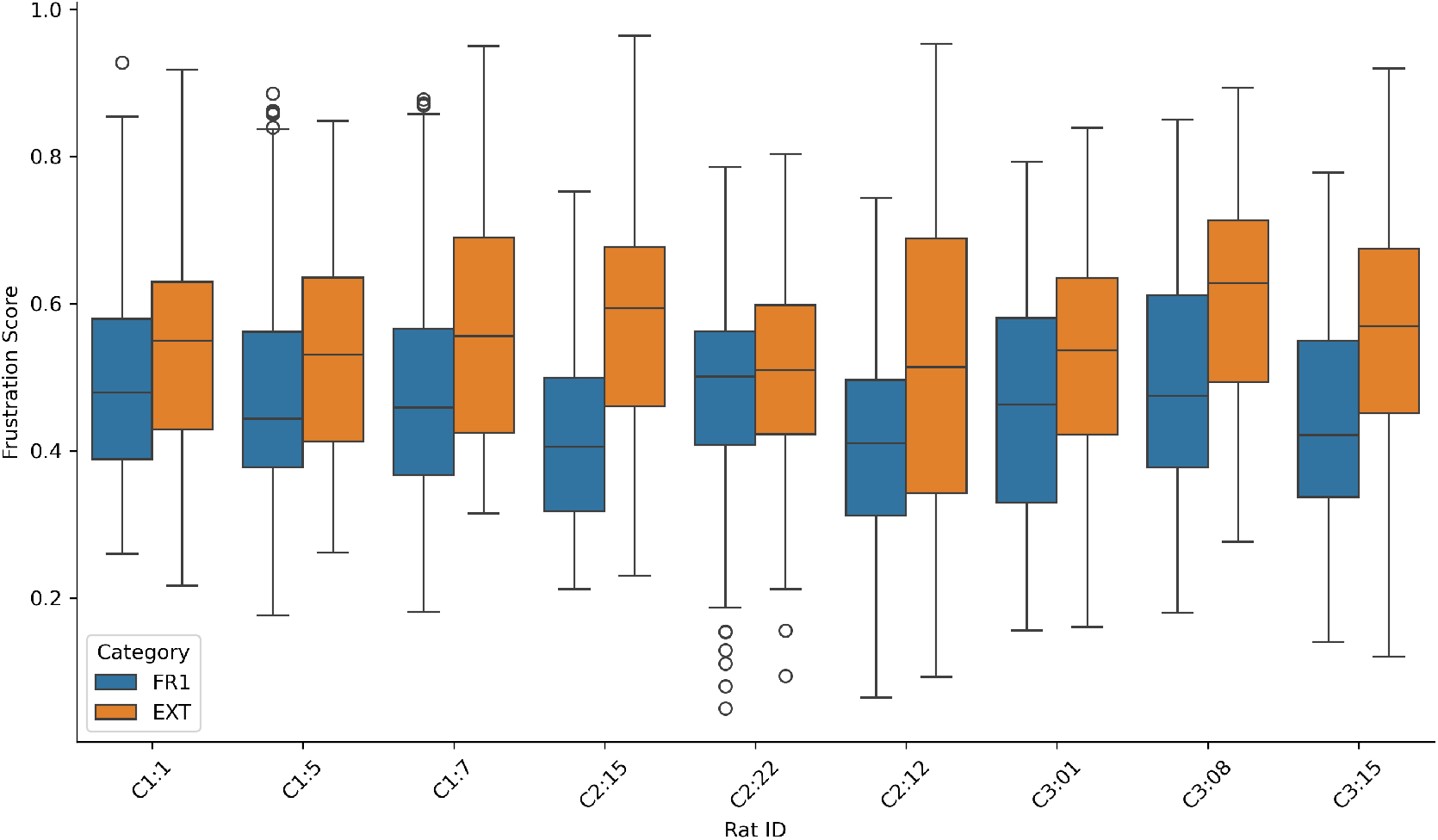
FR1 vs EXT frustration score in the holdout test set. The labels on the x-axis indicate the cohort and rat ID in the format C(cohort):(rat ID)

For inference, we visually examined three test rats under FR1 and PR settings in 7. A rolling average with window size of 10 was used to improve discriminability. The frustration score exhibited substantial between-animal variability. In rat 08, PR sessions showed consistently elevated frustration scores relative to FR1 sessions, whereas in others the separation was weaker and varied over the course of the session. Notably, all rats displayed temporal divergence between FR1 and PR trajectories as the session progressed.

## 4 Discussion

The results show several clear findings. First, barpress profiles can be used to draw inference about frustration-related behavior. Although each individual feature showed poor discriminability, with AUC values below 0.6, combining multiple features produced a much higher AUC in the GB model. Based on SHAP values, `Press Duration, Average Last 5 Samples, Mean Peak Sharpness,` and `Max Peak Width` were the most influential variables. In contrast, the AUC degradation curve showed that `Press Duration, Max Force, Average First 5 Samples, and Max Peak Width` contributed most to the model AUC. Together, the SHAP and permutation analyses suggest that model performance was driven by these six barpress features (`Press Duration, Average Last 5 Samples, Mean Peak Sharpness, Max Peak Width, Max Force, and Average First 5 Samples`).

It is important to note that `Press Duration and Max Force` are not isometric. Although one might expect that pressing a lever with greater force would take longer than pressing with less force, the correlation matrix in Figure 3A showed essentially no correlation between the two features (*r* = *−*0.07).

Thus, there was no force-time confound, and force-based features were able to contribute independently to model performance. More generally, the correlation structure suggested two distinct feature clusters. One consisting mainly of force-based features and the other consisting mainly of duration-based features.

One caveat from our previous work, which used barpress duration alone as a proxy for frustration state, was that not every barpress during extinction is clearly a “frustrated” barpress (i.e., one with a long duration)[6–9]. Even under frustration-related conditions, rats produced a mixture of long-duration barpresses and shorter, non-frustrated-looking barpresses, with the concentration of long barpresses increasing as frustration increased. Consistent with this idea, Figure 6 shows that although EXT sessions had a higher average frustration score than FR1 sessions, the majority of individual barpresses had similar frustration scores across conditions. Chunking consecutive barpresses and thus aggregating their frustration scores helped improve model discriminability. However, this benefit diminished as the chunk size became larger (e.g. chunk size *>*= 10). The increase in sensitivity from chunking provides additional opportunities for analysis. Within-session analyses would be best with no chunking or small chunks whereas animal-to-animal or session-to-session comparisons, such as research into individual differences would benefit from larger chunk sizes.

Our study does have limitations. First, the sesnsitivity of the model with no chunking was not stellar. Thus, this project can serve best as merely a proof-of-concept. Second, as shown in Figures 6 and 7, there was clear heterogeneity among rats in the inference PR data, meaning that not all rats exhibited the same frustration state at the same time. As a result, FR1 and PR barpresses could be difficult to distinguish at some points throughout the session; however, frustration scores in PR sessions increased at various points, which is consistent with frustration-realted behavior, and session-wide inference clearly showed increased frustration scores in the PR group vs. FR1. Heterogeneity among rats offers future potential in fine-tuning the model for specific rats for better sensitivity.

**Figure 7:**
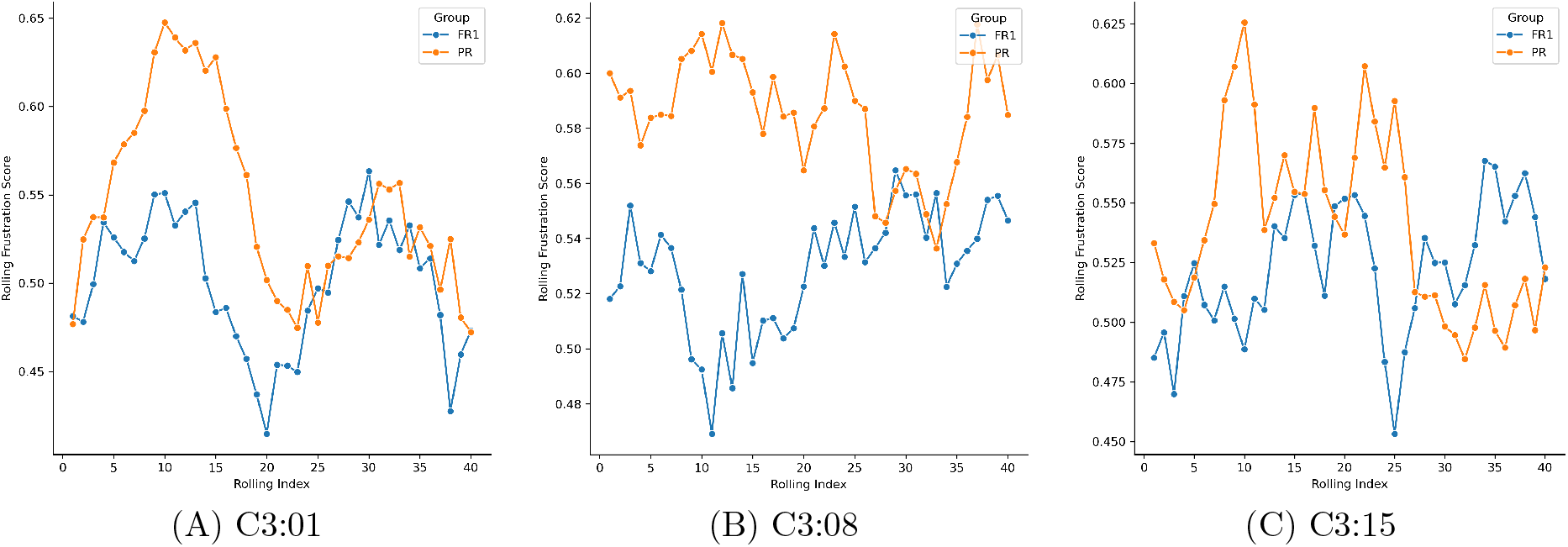
Predicted frustration score for the test rats. The cohort and rat ID in the format C(cohort):(rat ID)

Overall, our results demonstrate that machine learning models can adequately predict frustration state when applied to barpress profiles. These results show that assessing the force/time profile scores better than simple barpress durations alone. These results open the door to using barpress profiling to study other emotional or motivational states, such as “joy” in response to unexpected reward upshift, anticipation, or hunger motivation. This approach may also support empirical testing of frustration-based models of substance use disorders[3].

## Bibliography

[1] Green, T.A., Leibenluft, E., Li, Z., Ogawa, M., Sabariego, M., Mármol Contreras, Y., Bellaert, N., Papini, M.R.: Frustrative nonreward: A century of progress. Neurosci. Biobehav. Rev. 185(106651), 106651 (2026)

[2] Papini, M.R., Green, T.A., Mármol Contreras, Y., Torres, C., Ogawa, M., Li, Z.: Frustrative nonreward: Behavior, circuits, neurochemistry, and disorders. J. Neurosci. 44(40), 1021242024 (2024)

[3] Mármol Contreras, Y., Sanchez Rivas, R., Banh, S., Duncan, I.C., Earagolla, S., Green, T.A.: Frustration brake failure hypothesis of substance use disorders. Frontiers in Psychiatry 17, 1896594 (2026)

[4] Amsel, A., Roussel, J.: Motivational properties of frustration. i. effect on a running response of the addition of frustration to the motivational complex. J. Exp. Psychol. 43(5), 363–366 (1952)

[5] Jimènez-García, A.M., Ruiz-Leyva, L., Vázquez-Ágredos, A., Torres, C., Papini, M.R., Cendán, C.M., Morón, I.: Consummatory successive negative contrast in rats. Bio Protoc. 9(7), 3201 (2019)

[6] Vasquez, T.E., McAuley, R.J., Gupta, N.S., Koshy, S., Marmol-Contreras, Y., Green, T.A.: Lever-press duration as a measure of frustration in sucrose and drug reinforcement. Psychopharmacology 238(4), 959–968 (2021)

[7] Vasquez, T.E.S., Shah, P., Re, J.D., Laezza, F., Green, T.A.: Individual differences in frustrative nonreward behavior for sucrose in rats predict motivation for fentanyl under progressive ratio. eNeuro 8(5), 0136–212021 (2021)

[8] Mármol Contreras, Y., Vasquez, T.E.S., Shah, P., Payne, K., Di Re, J., Laezza, F., Green, T.A.: Bar press durations as a reliable and robust measure of frustration-related operant behavior: Sensitivity to incentive downshift and dose-response paradigms. PLoS One 18(12), 0296090 (2023)

[9] Mármol Contreras, Y., Dvorak, N.M., Tapia, C.M., Zaman, R., Annareddy, J., Balikosa, Y., Gupta, N.S., Rader, A.P., Vasquez, T.E.S., Koshy, S., Li, D., Balaji, V.K., Laezza, F., Green, T.A.: Depleting retinoic acid synthesis in the nucleus accumbens shell produces a protective phenotype for emotional reactivity and drug-taking in rats. Psychopharmacology (Berl.) 243(1), 59–74 (2026)

[10] Harris, M.B.: Aggressive reactions to a frustrating phone call. J. Soc. Psychol. 92(2), 193–198 (1974)

[11] Palomino, M., Cisneros-Plazola, M., Dubón, A., Pèrez-Trevinõ, V., López-Tolsa, G.E., Sosa, R.: Evidence for the extinction burst in grip force: A preregistered study (2026)

[12] Gèron, A.: Hands-On Machine Learning with Scikit-Learn, Keras, and TensorFlow: Concepts, Tools, and Techniques to Build Intelligent Systems, 3rd edn. O’Reilly Media, Sebastopol, CA (2022). pp. 54–56

[13] Lundberg, S.M., Lee, S.-I.: A unified approach to interpreting model predictions. arXiv preprint arXiv:1705.07874 (2017)

[14] Hastie, T., Tibshirani, R., Friedman, J.: The Elements of Statistical Learning: Data Mining, Inference, and Prediction, 2nd edn. Springer, New York, NY (2009). pp. 359–362

[15] BioRender: BioRender: Scientific Figure and Illustration Software. https://www.biorender.com/. Accessed August 21, 2026 (2026)

