## Supplemental Tables for "Detection of Frustration-related Operant Behavior in Rats via Machine Learning Methods"

Table S1: Description of Bar press features.

| Feature | Description |
|---|---|
| Max force | Maximum force (g) within a bar press. |
| Force variation rate | Maximum absolute differences between two consecutive force signals in the bar pr[illegible] |
| Average first 5 values | Mean of the first five force samples after press onset. |
| Average last 5 values | Mean of the last five force samples before press offset. |
| Peak sharpness | peak's prominence divided by its width at half-prominence. |
| Skewness | Skewness of the force samples within a press. |
| Kurtosis | Kurtosis of the force samples within a press. |
| Press duration | Total time (s) the force trace stays above the press threshold. |
| Peak Number | Count of detected peaks within a press. |
| Peak width | Duration (s) associated with detected peaks within a press. |

Table S2: hyperparameters in `find_peaks`

| Hyperparameter | Definition | Values searched | Optimal value |
|---|---|---|---|
| prominence | How much a peak stands out from neighboring samples. | {0.2, 0.5, 1, 2, 5, 10} | 10 |
| height | Minimum required peak amplitude. | {1, 2, 5, 10, 20, 50} | 20 |
| distance | Minimum number of samples between adjacent peaks. | {1, 3, 5, 10, 20, 50} | 10 |
| width | Minimum peak width. | {1, 2, 3, 5, 10, 20} | 5 |

Table S3: Hyperparameter search space and selected values for the gradient boosting model.

| Hyperparameter | Values Searched | Selected Value |
|---|---|---|
| max_depth | $\{1, 3, 5, 7, 9\}$ | 3 |
| subsample | $\{0.6, 0.75, 0.9, 1.0\}$ | 1.0 |
| colsample_bytree | $\{0.6, 0.75, 0.9, 1.0\}$ | 0.6 |
| reg_lambda | $\{0, 0.1, 0.5, 1, 3, 10\}$ | 0.1 |
